# Current genomic deep learning architectures generalize across grass species but not alleles

**DOI:** 10.1101/2024.04.11.589024

**Authors:** Travis Wrightsman, Taylor H. Ferebee, M. Cinta Romay, Arun S. Seetharam, Taylor AuBuchon-Elder, Alyssa R. Phillips, Michael Syring, Matthew B. Hufford, Elizabeth A. Kellogg, Edward S. Buckler

## Abstract

Non-coding regions of the genome are just as important as coding regions for understanding the mapping from genotype to phenotype. Interpreting deep learning models trained on RNA-seq is an emerging method to highlight functional sites within non-coding regions. Most of the work on RNA abundance models has been done within humans and mice, with little attention paid to plants. Here, we benchmark four genomic deep learning model architectures with genomes and RNA-seq data from 18 species closely related to maize and sorghum within the Andropogoneae. The Andropogoneae are a tribe of C4 grasses that have adapted to a wide range of environments worldwide since diverging 18 million years ago. Hundreds of millions of years of evolution across these species has produced a large, diverse pool of training alleles across species sharing a common physiology. As model input, we extracted 1,026 base pairs upstream of each gene’s translation start site. We held out maize as our test set and two closely related species as our validation set, training each architecture on the remaining Andropogoneae genomes. Within a panel of 26 maize lines, all architectures predict expression across genes moderately well but poorly across alleles. DanQ consistently ranked highest or second highest among all architectures yet performance was generally very similar across architectures despite orders of magnitude differences in size. This suggests that state-of-the-art supervised genomic deep learning models are able to generalize moderately well across related species but not sensitively separate alleles within species, the latter of which agrees with recent work within humans. We are releasing the preprocessed data and code for this work as a community benchmark to evaluate new architectures on our across-species and across-allele tasks.

## Introduction

Non-coding regions of the genome are well-known to be as important as coding regions for understanding how genotype determines phenotype (Finucane et al., 2015; Rodgers-Melnick et al., 2016). Though tools like AlphaFold2 (Jumper et al., 2021) have dramatically improved our ability to study coding sequence, similarly performing tools do not yet exist for non-coding regions. Nevertheless, over the last decade deep learning models have rapidly improved performance in predicting non-coding genomic features such as chromatin accessibility (Kelley, 2020; Wrightsman et al., 2022), transcription factor binding (Žiga Avsec, Weilert, et al., 2021; Mejía-Guerra & Buckler, 2019), and RNA abundance (Žiga Avsec, Agarwal, et al., 2021; Linder et al., 2023) directly from DNA sequence. These models can then be queried to highlight functional non-coding sites, which can be useful for filtering large sets of variants down to promising genome editing targets. Further, since most of the modeling work has been done on human and mouse data, there is a need to benchmark their performance in plants.

Models that predict RNA abundance from sequence are particularly attractive due to the relatively cheap cost and standardized protocols of RNA-seq. However, there is room for improvement in these models across a number of areas. While RNA abundance models have shown high performance across genes, recent work in humans (Huang et al., 2023) has highlighted their lack of sensitivity across individuals. Some expression model architectures (Žiga Avsec, Agarwal, et al., 2021; Linder et al., 2023) include coding sequence in the input, which is known to lead to overfitting on gene family instead of true regulatory sequence differences (Washburn et al., 2019). There is also a tendency to maximize data when training these models, without actually measuring the rate of diminishing returns for each additional observation. Finally, while multiple species have been included in some training sets, it is common to test on a set of held-out chromosomes within the training species, rather than testing on a completely held-out species.

Deep learning models benefit from large and diverse training sets of different tissues and genotypes, which are rarely available outside model species. To train RNA expression models on larger sample sizes, we leveraged new long-read genomes and RNA-seq data from 15 wild species of the Andropogoneae tribe. Diverging around 17.5 million years ago (Welker et al., 2020), the Andropogoneae includes globally staple crop plants such as maize, sorghum, and sugarcane. Millions of years of evolution within the tribe has provided a large, diverse pool of training alleles. Sorghum and maize diverged around 12 million years ago (Mya), on the order of the human-chimpanzee split (6–10 Mya), but have a 10-fold higher rate of nucleotide divergence (Chimpanzee Sequencing and Analysis Consortium, 2005; Zhang et al., 2017).

We tested four sequenced-based genomic deep learning architectures, DanQ (Quang & Xie, 2016), HyenaDNA (Nguyen et al., 2023), FNetCompression (Pipoli et al., 2023), and a smaller version of Enformer (Žiga Avsec, Agarwal, et al., 2021), on their ability to predict across species and alleles. DanQ is one of the earliest genomic deep learning architectures, leveraging a long short-term memory recurrent layer to learn the syntax and grammar of motifs detected by a convolutional layer. Enformer is a massive transformer architecture with a context size near 100 kilobases that is among the best performing models for human expression prediction. HyenaDNA is a novel architecture capable of handling long context windows of up to a million base pairs. FNetCompression combines a fast Fourier transform with multi-head attention to efficiently learn from sequences of up to 10 kilobases with a few orders of magnitude less parameters than the other architectures.

We aimed to investigate, from a plant perspective, two major open questions in expression modeling from sequence: 1) How well do current sequence-based deep learning architectures generalize across species? and 2) How sensitive are these models across individuals?

## Results

### Current genomic deep learning architectures generalize across species

We trained all four architectures on genomic sequence and RNA-seq data from 15 species within the Andropogoneae clade (Figure 1a). Our validation set consisted of the two sampled species closest to *Zea mays, Tripsacum zopilotense* and *Zea diploperennis*, all three of which fall within the Tripsacinae subtribe that diverged a few (0.6–4) million years ago (Chen et al., 2022; Welker et al., 2020). Our test set was the 26 inbred parents of the maize NAM population (Hufford et al., 2021; Yu et al., 2008), held out until hyperparameters were frozen. As input, we extracted 1,026 base pairs upstream of the translation start site to match HyenaDNA’s “tiny” configuration (Figure 1b). We trained and evaluated all architectures on two regression tasks, maximum expression across tissues and absolute expression in leaf, as well as two classification tasks, expressed in any tissue and expressed in leaf (Figure 1b).

**Figure 1.**
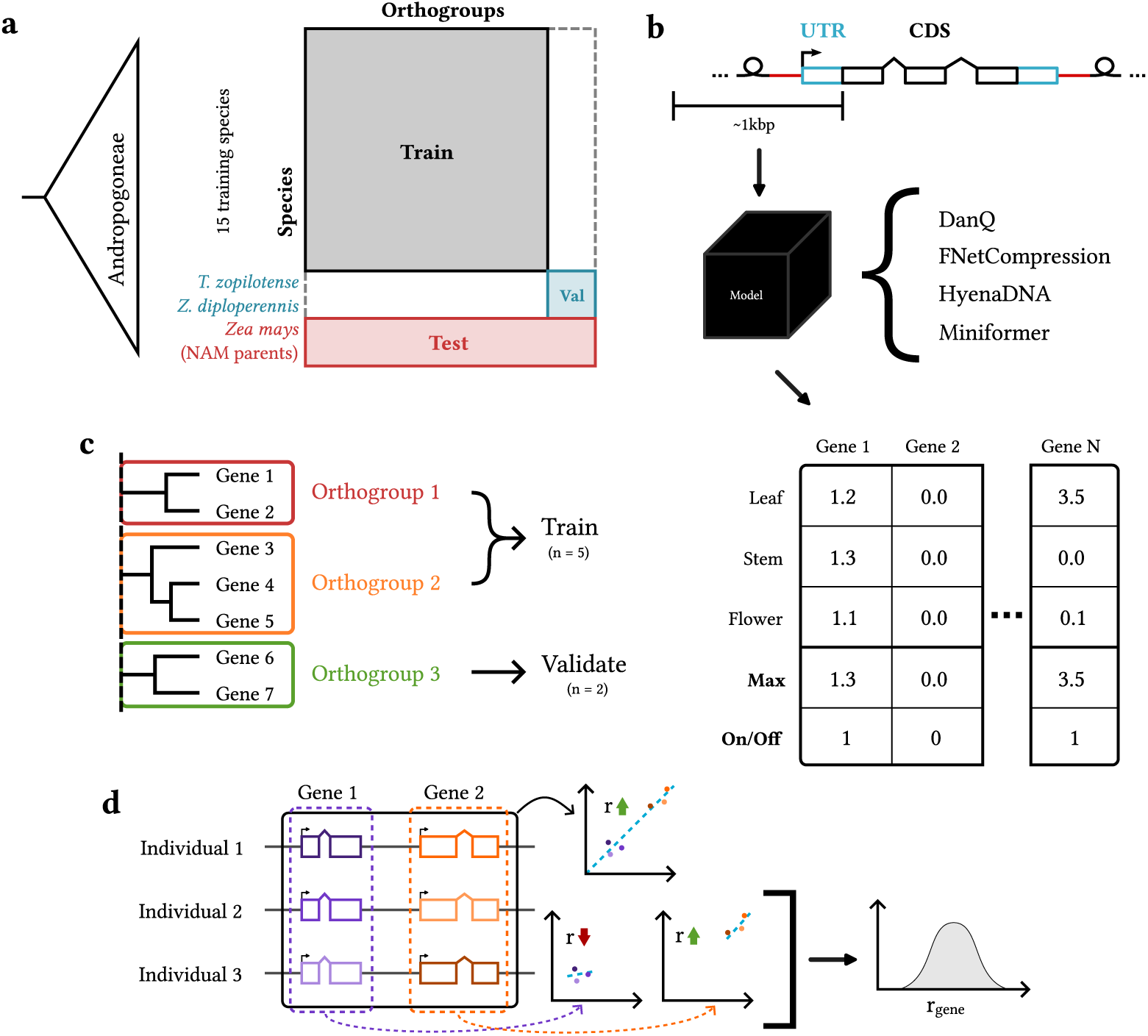
Methods overview: a) data splits; b) models, features, and targets; c) orthogroup-guided splitting; d) metrics (across-vs. within-gene performance)

Though benchmarking of sequence-based models has been done within humans (Karollus et al., 2023) and across species in the training set (Kelley, 2020; Levy et al., 2022), there has been little evaluation on entirely held out species. To establish a baseline in plants, we measured performance of all architectures, tasks, genes, and data splits (Figure 2a–d). Rankings by Spearman correlation on the test set are inconsistent, except that DanQ performed the best or tied closely with the best across all tasks. Remarkably, DanQ performs only slightly worse (Figure 2a; Δ*r* = 0.09) than Enformer in a recent within-human single tissue benchmark (Huang et al., 2023) despite predicting on an unobserved species. Despite having moderate Spearman and Pearson correlation (Supplemental Figure 1), DanQ’s predictions on the test set are still underwhelming (Supplemental Figure 2). We observed test set auROC scores in the any expression task slightly lower than previous results (Washburn et al., 2019) on promoter expression classification models trained and tested only within maize. Taken together, these results support modern genomic deep learning architectures are capable of generalizing almost as well across closely related species as they do within species.

**Figure 2.**
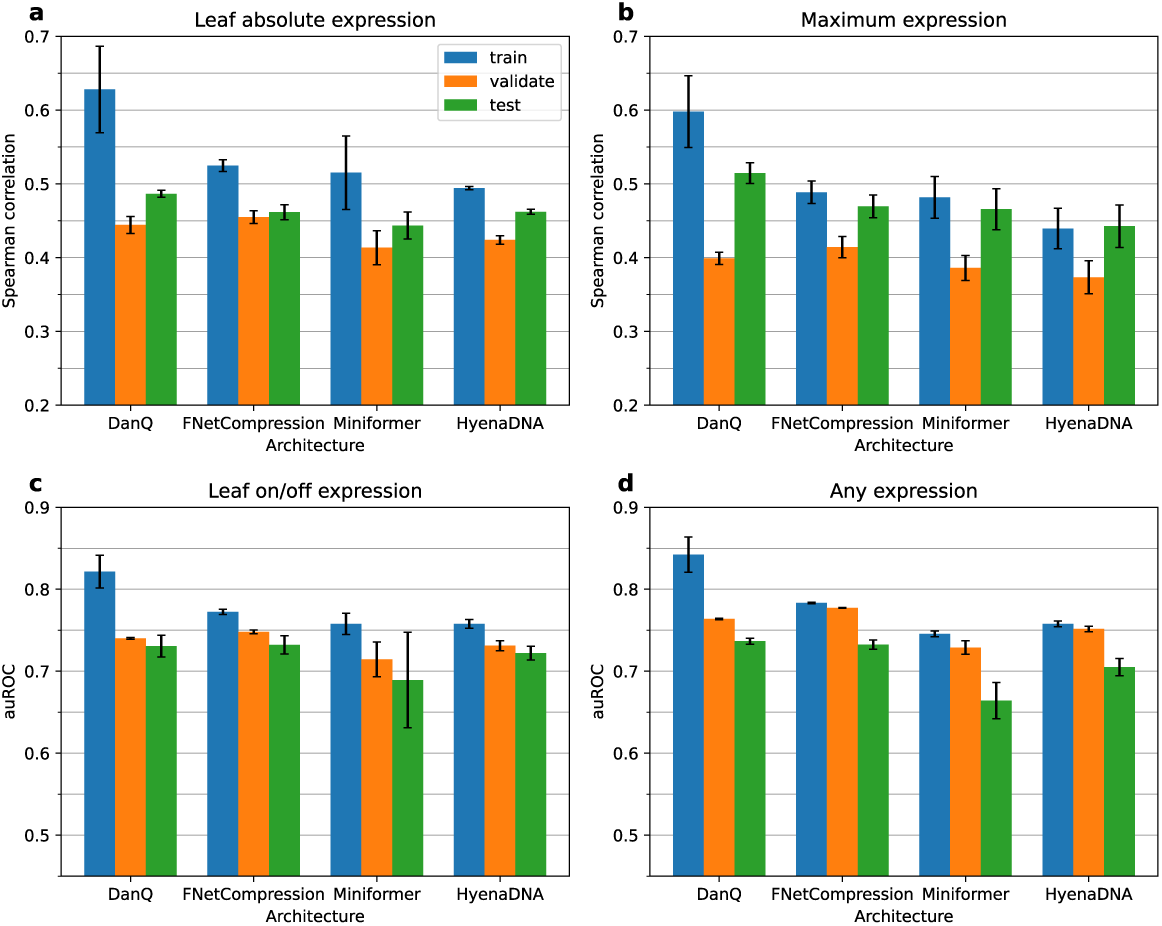
a–d) Model performance across all genes and data splits. Each subfigure shows the performance of all architectures on a single task. Error bars represent one standard deviation from the mean in each direction.

### Data quantity matters more than composition for modeling RNA abundance across species

Despite the growing number of plant genomes with transcriptomic data (Sreedasyam et al., 2023), each genome added to the training set increases training time and may give diminishing returns. We measured changes in DanQ’s performance on progressively larger fractions of the training data, iteratively adding sequences from a set of genomes or randomly from across all training genomes. Pearson correlation on the validation set rises until approximately 200,000 observations when it begins to show diminishing returns for larger training set sizes (Figure 3a). However, the slope remains positive between the half size and full size data points, suggesting room for improvement with further observations. Comparing iteratively adding whole genomes to randomly sampling an equivalent number of alleles from the entire training set, there are only substantial differences when using less than 8 genomes, with random performing worse than 4 whole genomes. The ablation results clearly support the use of further data to achieve higher performance across genes, which can come from sequencing additional related species.

**Figure 3.**
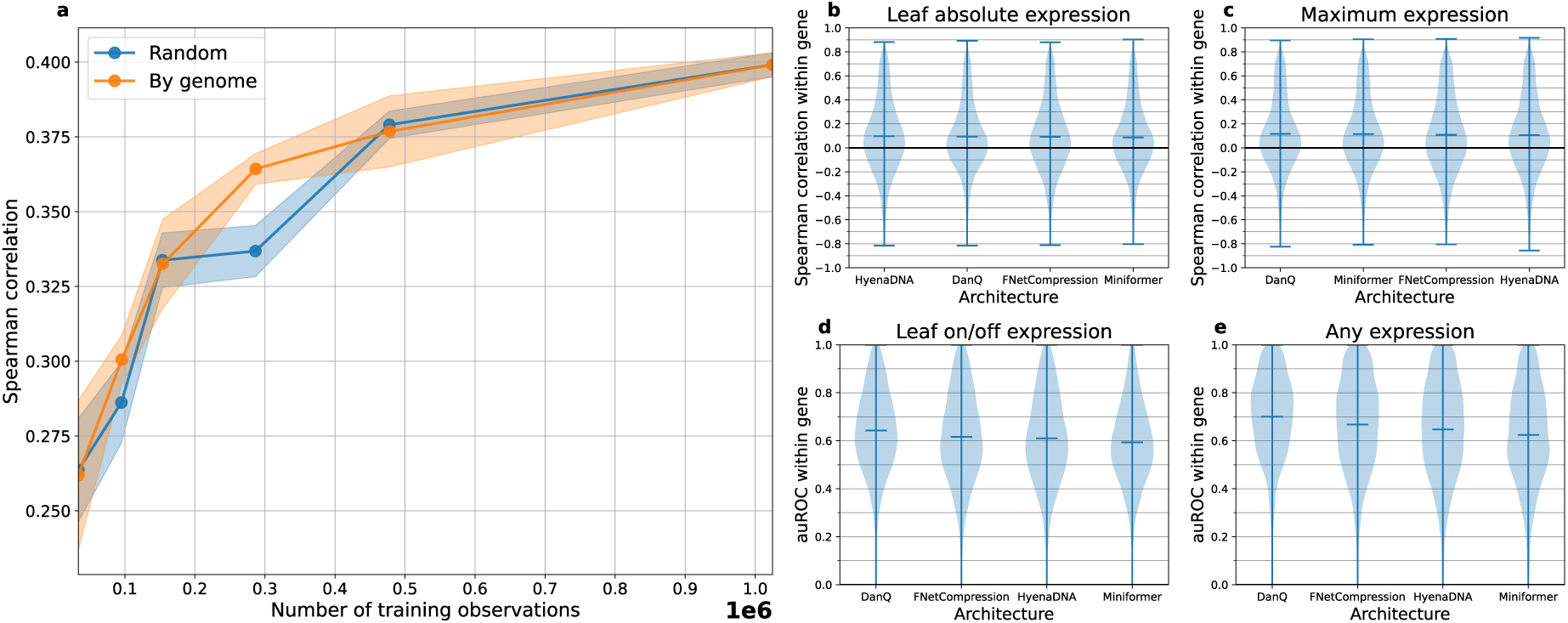
a) Validation set performance of DanQ on the maximum expression task across varying training set sizes and compositions. Points on the lines are mean Pearson correlation across replicate training runs. The standard deviation across replicates is shaded around the mean line. The exponent scale of the bottom axis is denoted in bold on the right. b–e) Distributions of model performance within orthogroups for each task. Architectures are sorted from highest (left) average within-orthogroup performance to lowest (right). Bars within the violins represent the mean of the distribution.

### Current architectures poorly generalize across individuals of an inbred maize panel

Recent work (Huang et al., 2023) has shown that current models poorly explain expression variation across individuals. Since our test set is a collection of maize alleles with an order of magnitude more diversity than humans (Chia et al., 2012), we looked at the distribution of test set performance within each orthogroup and expected to see similarly low or even lower performance. We only considered orthogroups that had at least 20 orthologs to have sufficient sample size for calculating correlation or auROC. We saw much lower average within-orthogroup Spearman and Pearson correlations as well as auROC compared to the global across-gene metrics, except for the any expression task (Figure 3b–e; Supplemental Figure 3), which also shows clear differences between architectures. The average within-orthogroup Spearman correlation in the single tissue regression task is double (*r* = 0.092) than what was observed with Enformer (*r* = 0.045) in humans.

As an alternative allele-level comparison, we also looked at how well DanQ predicted expression differences between all pairs of maize ortholog promoter alleles within a orthogroup. We observed a general positive relationship between the two, but there is still quite a bit of noise in the predictions (Figure 4, left). The Pearson and Spearman correlation coefficients between the observed and predicted fold changes were only 0.22 and 0.08, respectively. Strikingly, pairs of orthogroups that are two orders of magnitude apart in expression level are still sometimes predicted with the incorrect direction. Despite current architectures showing promising across gene performance in unobserved species, they still struggle across shorter evolutionary timescales, similar to what was seen in humans (Huang et al., 2023).

**Figure 4.**
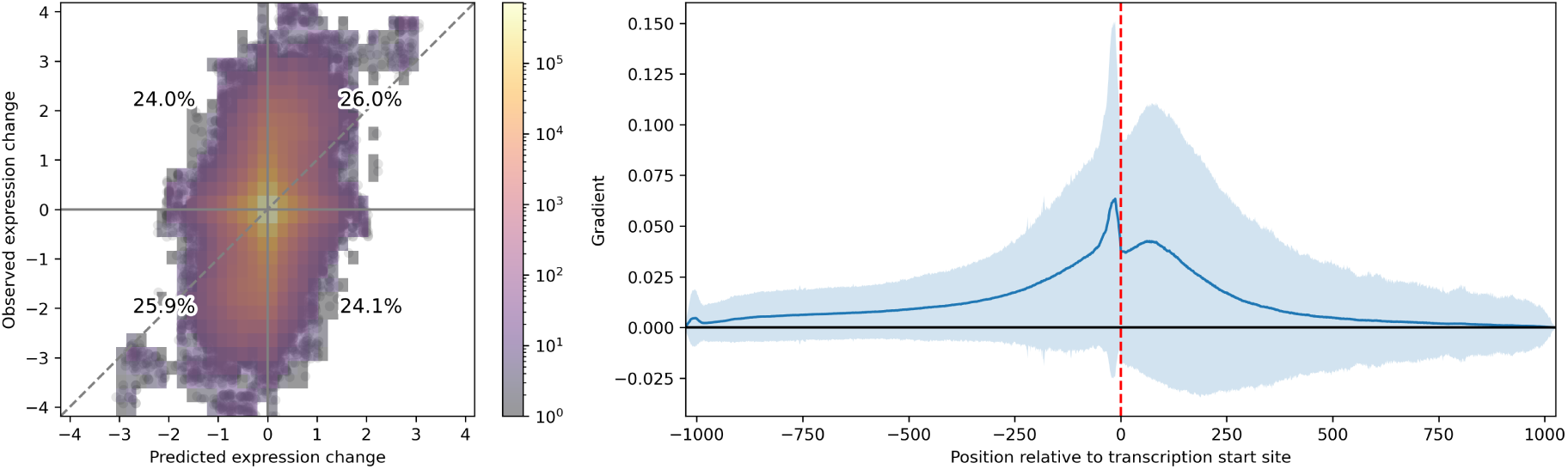
Left: Predicted versus observed log_10_ expression change in leaf between all NAM ortholog promoter allele pairs within an orthogroup. Percentages in the middle of each quadrant display the proportion of non-zero data points in that quadrant. Right: Saliency map for DanQ trained on maximum expression task. The mean across all B73 genes is plotted as a line, with a single standard deviation shaded above and below.

### The maximum expression regression model focuses on the core promoter region

Based on theory and prior interpretation work on expression models (Mendoza-Revilla et al., 2023), we hypothesized our expression models would also pay most attention to the region surrounding the transcription start site. Looking at the average saliency map for DanQ across all B73 genes on the maximum expression task we see that DanQ indeed focuses on the core promoter region and the 5′ UTR (Figure 4, right). There is relatively high variance in the gradient around the transcription start site, taping off with increasing distance, though decaying slower in the UTR than promoter. This hyperfocus on the core promoter region could be why DanQ and other architectures struggle to distinguish expression differences in highly related sequences, since functional mutations are less likely to accumulate in this highly constrained region.

## Discussion

Here we have shown that four genomic deep learning architectures are capable of generalizing across species, though they also show the same lack of allelic sensitivity seen in humans (Huang et al., 2023). FNetCompression’s performance is particularly remarkable because it has several orders of magnitude fewer parameters than DanQ (57k versus 1.6m, respectively). Large foundation models such as AgroNT (Mendoza-Revilla et al., 2023) show promising results within the training species, but FNetCompression suggests smaller, more efficient models, perhaps also utilizing a fast Fourier transform, are worth further exploration. Since the Pearson correlations we observed are still far from perfect, it is worthwhile to note that we do not expect *cis* sequence-based models to ever reach perfect correlation, as *cis* factors explain only a third of the genetic variation in expression in maize (Giri et al., 2021). The fact that our models show across individual performance in maize (*r*_*s*_ = 0.092) double that observed in humans (Huang et al., 2023) is puzzling. Population genetics has shown that maize has an order of magnitude more genetic variance than humans (Chia et al., 2012), yet our models are generalizing across maize individuals better than what was observed across human individuals. Unlike our validation set, our maize test set includes orthologs of sequences in our training set, which may result in slightly inflated performance estimates. However, this inflation is expected to be less than when coding sequences are included in the model (Washburn et al., 2019), as was the case for the human benchmark. More work will be needed to investigate this and, more generally, where the remaining errors are being made in these models.

This stringent benchmark, both across species and across individuals in a held-out species, is something that all expression models should be continually evaluated against to get a better sense of generalizability than within species testing. Our ablation results show that we are not yet saturated in terms of training data, meaning there is a need for further benchmarks on larger sets of species. Training across species presents the opportunity to not only leverage larger data sets but to learn the general rules of eukaryotic gene expression. Future work should consider training across many distantly-related species to learn general rules, then successively fine-tuning within clades to learn lineage-specific patterns. Consideration of data balance may be necessary, as prevalent polyploidy within plants (Song et al., 2023) leads to vastly different gene counts across species and complicates transcript quantification. However, scaling to bigger data will come with metadata challenges, exacerbated by the plethora of standards across databases (McQuilton et al., 2016). While considering better architectures with higher data needs, it will be increasingly important to better organize expression databases. Lastly, with the rising utilization of foundation models trained across massive numbers of genomes, it will also be critical to maintain true hold-out species for fair model evaluation. A CASP-like competition for RNA abundance modeling may be useful for this, as new sequence-based models of non-coding biology are developed.

## Materials and Methods

The companion Zenodo repository (Wrightsman et al., 2024) contains the source code required to reproduce this manuscript. Pandas (McKinney, 2010) and Polars (Vink et al., 2023) were used to process tabular data. Matplotlib (Hunter, 2007) was used for plotting figures. GNU parallel (Tange, 2024) was used for managing parallel execution of some analyses. This manuscript is written in and rendered using Typst (Haug, 2022; Mädje, 2022).

### Software Environments

Software environments were managed with pixi (Arts et al., n.d.). Packages were downloaded from the conda-forge (Conda-Forge Community, 2015) and Bioconda (Grüning et al., 2018) Conda channels. The exact software versions used in this work are defined in configuration files within the manuscript’s companion repository.

### Data preprocessing

All genome assemblies and annotations were downloaded from MaizeGDB (Hufford et al., 2021; Portwood et al., 2018). Version 5 of the B73 assembly and annotation was used. For PanAnd, version 1 of the assemblies and version 2 of the annotations were used. B73 and other NAM parent RNA-seq data were downloaded from ArrayExpress accessions E-MTAB-8628 and E-MTAB-8633, respectively. Other Andropogoneae RNA-seq data were downloaded from NCBI accession PRJNA1098707. Transcript quantifications were obtained using quantify-RNA-pipeline (Wrightsman, 2023). RNA-seq samples with less than 5 million mapped reads were dropped from further analysis. eggnog-mapper (Cantalapiedra et al., 2021) was used to assign proteins to Poales orthogroups.

*Zea mays* genes were assigned to the test set. 90% of orthogroups were randomly chosen as training orthogroups, with the other 10% used for validation. *Zea diploperennis* and *Tripsacum zopliotense* genes in the validation orthogroups were used as the validation set. Genes in the training orthogroups in all remaining Andropogoneae genomes were assigned to the training set.

Annotations were processed using gffutils (Dale, 2023). For each gene, the highest expressed transcript across all tissues was selected as a representative gene model. TPM values from other transcripts of the same gene that share the same transcription start site were added to the chosen transcript’s TPM. For the purposes of computing max expression, only leaf, shoot, and floral tissues were used as only those tissues had sufficient sampling across all species. The any tissue and leaf on/off expression classification task targets were binarized from the max expression and leaf absolute expression regression task targets (TPM). Specifically, TPM values of zero were kept as zero (unexpressed) and TPM values greater than zero were set to one (expressed).

### Model architectures

Exact hyperparameter settings for each architecture is specified in configuration files within the companion repository. DanQ (Quang & Xie, 2016) and FNetCompression (Pipoli et al., 2023) were both converted to PyTorch (Paszke et al., 2019), keeping all hyperparameters identical. Miniformer is a scaled-down version of the Enformer (Žiga Avsec, Agarwal, et al., 2021) architecture, with lower model dimensions and fewer layers. HyenaDNA (Nguyen et al., 2023) was used in the “tiny” configuration. Classification architectures were identical to their regression counterparts except that the final activation function was changed to sigmoid.

### Training

PyTorch Lightning (Falcon & The PyTorch Lightning team, 2019) and Hydra (Yadan, 2019) were used to orchestrate the training process and provide an interface for the data loader. MLFlow (Zaharia et al., 2018) was used to track experiment parameters and metrics as well as store model artifacts. As input, 1,026 base pairs upstream from the translation start site were extracted. The sequence was reverse complemented if the transcript was on the negative strand. If the model used 1-hot encoded sequence as input, the sequence was 1-hot encoded and then padded or trimmed as needed to be exactly 1,026 base pairs in length. If the model used tokens as input, the input sequence was tokenized to a max length of 1,026, padding as needed. If the task was regression (max or absolute leaf expression), TPM values were log_10_(TPM + 1) transformed. Training continued until the validation loss failed to decrease after three epochs. The model checkpoint at the end of the epoch with the lowest validation loss was kept. Each combination of architecture and task was run three times with different initial seeds to estimate model robustness. Any runs that failed to converge were restarted with a different seed value until a total of three converged runs were obtained.

### Ablation

“By genome” ablation was performed by filtering to one, two, three, four, or eight training genomes, in order of increasing phylogenetic distance from the test set. The first eight genomes, in order, were *Tripsacum dactyloides* “FL_9056069_6”, “McKain_334-5”, *Elionurus tripsacoides, Hemarthria compressa, Thelepogon elegans, Sorghastrum nutans, Ischaemum rugosum*, and *Pogonatherum paniceum*. “Random” ablation randomly sampled an equivalent number of observations from the total set of observations from all 15 training species. For example, if the “By genome” ablation had two genomes with 30,000 and 35,000 observations, then the corresponding “random” ablation experiment would randomly sample 65,000 observations from the total training set. Each ablation run was repeated six times to measure robustness.

### Ortholog contrast

The first DanQ training run model was used to predict the expression for each transcript in the test set. All possible pairs of orthologs within each orthogroup were generated for the contrast. Orthogrups were filtered to those that contained between 20 and 35 members, to avoid private genes and retroelements and have sufficient sample sizes to calculate correlation and auROC.

### Saliency map

Captum (Kokhlikyan et al., 2020) was used to compute saliency. For each position, the absolute value of saliency for each channel was summed. The mean and standard deviation of this sum was computed across all B73 genes.

## Acknowledgements

This work was funded by the USDA-ARS and NSF PanAnd grant (Award 1822330). T.H.F. was supported by a USDA NIFA AFRI predoctoral fellowship (TF: 2022-67011-36564). Some compute resources were provided by the Cornell University BRC Bioinformatics Core Facility (RRID:SCR_021757). This research used resources provided by the SCINet project and/or the AI Center of Excellence of the USDA Agricultural Research Service, ARS project numbers 0201-88888-003-000D and 0201-88888-002-000D. Aaron Gokaslan helped design and debug the data loader.

## Author Contributions

In terms of the Contributor Roles Taxonomy (CRediT): All authors were responsible for Writing – review & editing. T.W. was additionally responsible for Conceptualization, Data curation, Formal analysis, Investigation, Methodology, Project administration, Software, Validation, Visualization, and Writing – original draft. T.H.F. for Conceptualization, Data curation, and Methodology. M.C.R. for Data curation and Project administration. A.S., T.A., A.R.P., M.S., and M.B.H. for Resources. E.A.K. for Data curation and Resources. E.S.B. for Conceptualization, Funding acquisition, Methodology, Project administration, and Supervision.

## Supplemental Material

**Supplemental Figure 1:**
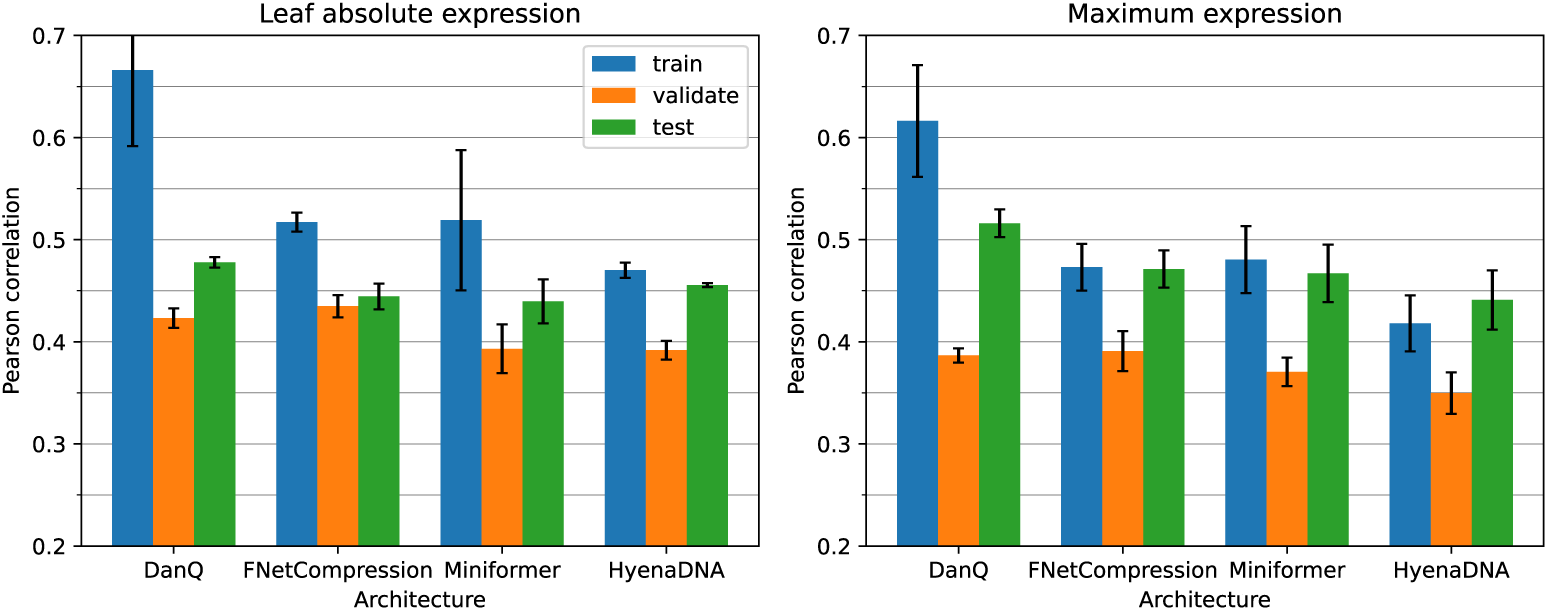
Pearson correlation in the regression tasks across all genes and data splits. Each plot shows the performance of all architectures on a single task. Error bars represent one standard deviation from the mean in each direction.

**Supplemental Figure 2:**
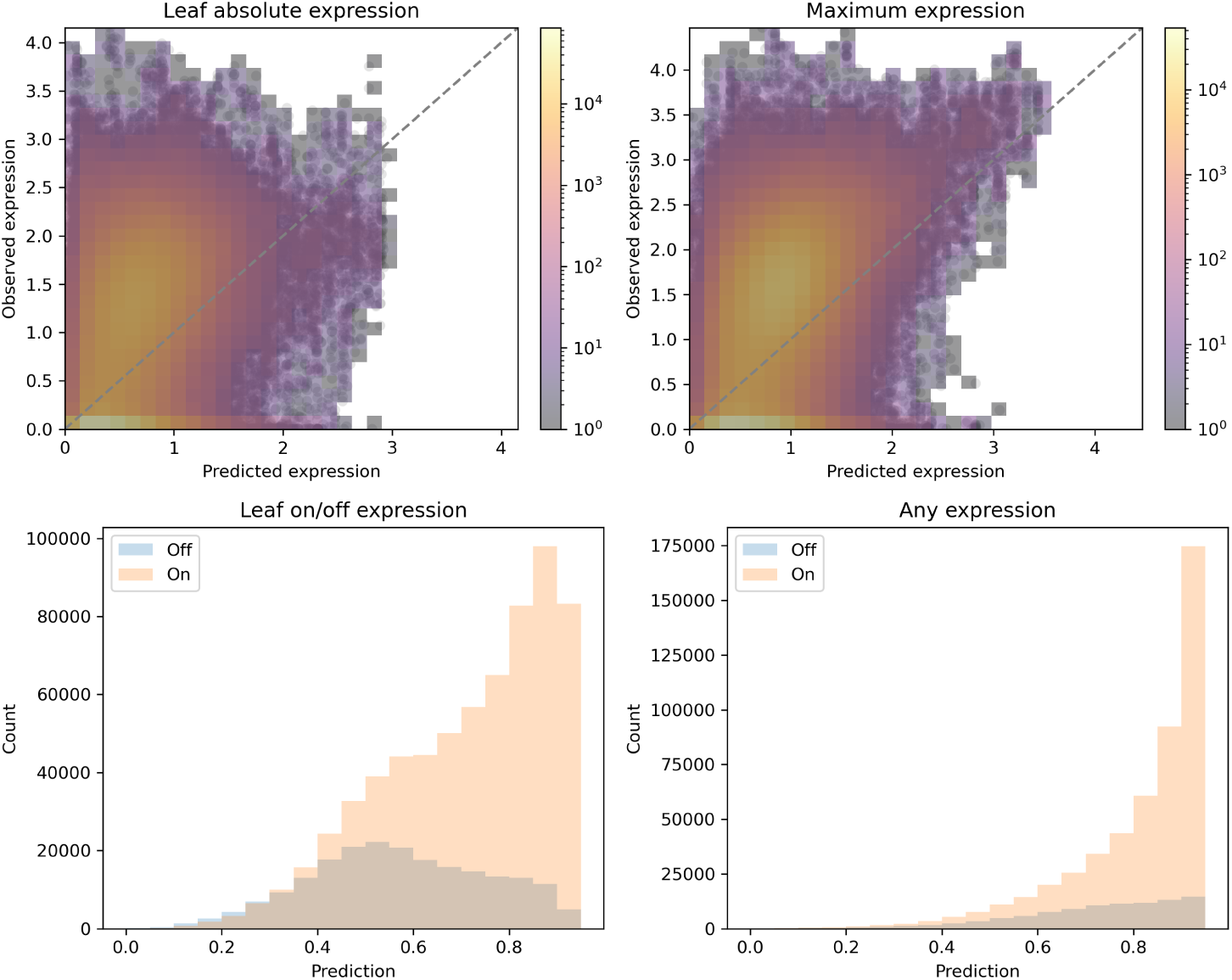
DanQ predictions on the test set across all tasks. The model from the first training run was used for predictions. Regression tasks (top) are on the log scale. Color in the regression task histogram scatterplots represents the number of observations within that area.

**Supplemental Figure 3:**
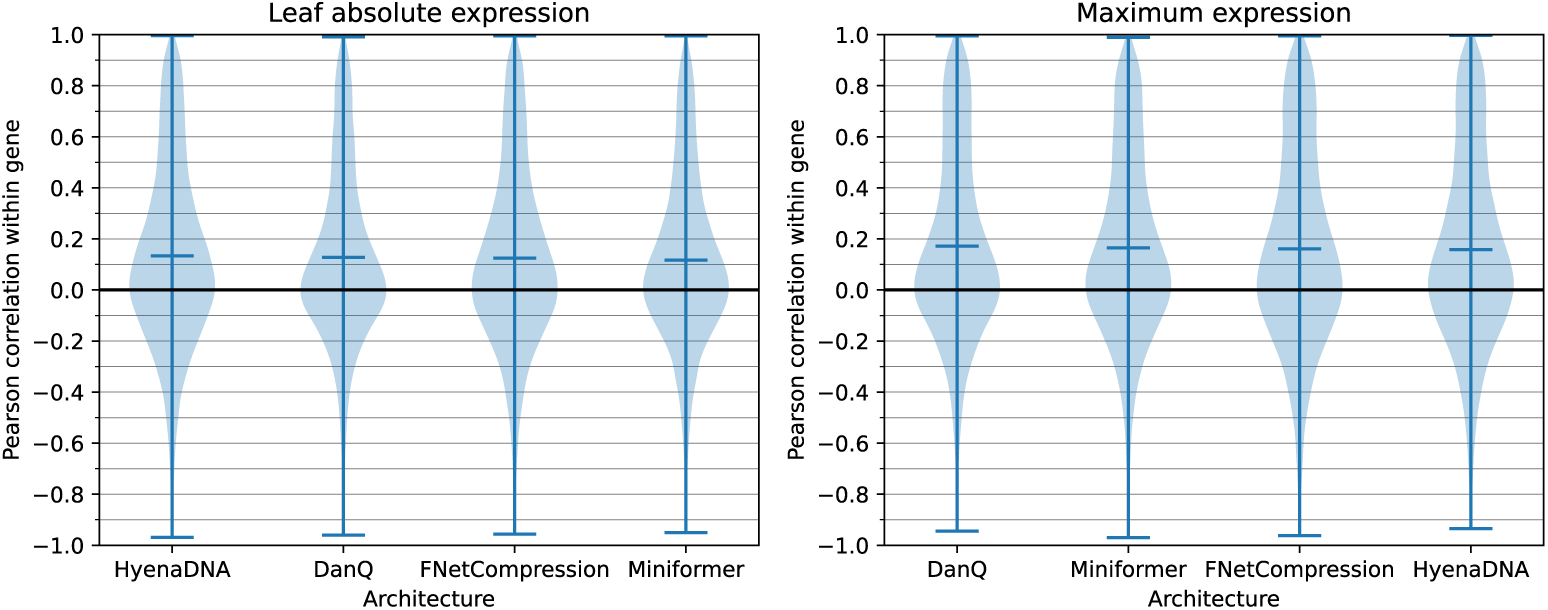
Distributions of Pearson correlation within orthogroups for each task. Architectures are sorted from highest (left) average within-orthogroup Pearson correlation to lowest (right). Bars within the violins represent the mean of the distribution.

